# Rewiring V-type and K-type enzyme allostery through subunit interface mutations

**DOI:** 10.64898/2026.05.29.728626

**Authors:** Apala Chaudhuri, Federica Maschietto, Olivia Enny, Victor S. Batista, J. Patrick Loria

## Abstract

Allosteric regulation in the heterodimeric enzyme IGPS depends on long-range communication between the effector-binding HisF subunit and the catalytic HisH subunit. This signaling occurs through a densely packed interdomain interface enriched in conserved noncovalent contacts. Here, we use targeted interface mutations to determine *how* specific interfacial contacts tune the structural and dynamic features that govern allosteric control. The hK181A variant, which disrupts a critical salt bridge with fD98, converts IGPS into a constitutively more active enzyme, increasing basal glutaminase activity and substrate affinity. By contrast, hR18A, which disrupts a secondary salt bridge with fE71, weakens effector-induced activation, revealing functional asymmetry among interfacial interactions. NMR chemical shift perturbation and CPMG relaxation dispersion experiments show that hK181A remodels millisecond-timescale dynamics throughout HisF, consistent with molecular dynamics simulations indicating enhanced sampling of catalytically competent conformations. Network traffic analysis of correlated communication pathways, combined with energetic analysis, further shows that enthalpic and entropic contributions are redistributed to rewire long-range allosteric signaling. Together, these results identify specific interfacial residues as molecular gates that shape the conformational ensemble accessible to IGPS and show how interface reengineering can be used to rationally reprogram allosteric output.

**Significance Statement:** Enzymes often work like molecular switches: binding at one site can change activity at another distant site. How to predict or redesign this communication remains a major challenge. Using imidazole glycerol phosphate synthase (IGPS), we show that changing single amino acids at the interface between its two protein subunits can alter how the enzyme responds to regulation. One substitution shifts IGPS toward a response that changes both catalytic rate and substrate binding, whereas another weakens activation by disrupting communication across the interface. These findings identify interfacial residues that act as control points in an allosteric network and suggest a practical strategy for engineering enzymes with customized regulatory behavior.

---

**A**llostery provides a general mechanism by which ligand binding at one site modulates protein function at a distant site.(1, 2) In enzymes, allosteric ligands can alter the maximal catalytic rate (V-type allostery), the apparent substrate affinity (K-type allostery), or both.(3) These regulatory outputs arise from ligand-dependent shifts in conformational ensembles that contain states of differing catalytic competence.(4) Defining how specific structural elements bias these ensembles remains central to understanding, and ultimately engineering, enzyme regulation.

Imidazole glycerol phosphate synthase (IGPS) is a well-established model for long-range allosteric communication.(5) IGPS is a heterodimer composed of glutaminase subunit HisH and the cyclase subunit HisF. Catalysis requires tight coupling between glutamine hydrolysis in HisH and product formation in HisF. Effector binding in HisF (e.g., PRFAR or IGP), activates glutamine hydrolysis in HisH at a site more than 25 Å away from the HisF active site (Fig. 1a).(6, 7) This activation is associated with changes in characteristic dynamic motions, including altered mobility of HisF *loop1* near the effector-binding site.(8–11) These motions propagate across the subunit interface and promote a catalytically competent HisH active site. A key event is disruption of the inhibitory hydrogen bond between the carbonyl of hP10 and the amide of hG50, which permits formation of the *oxyanion hole* required for glutamine hydrolysis (Fig. 1b and c).(5, 12)

**Fig. 1.**
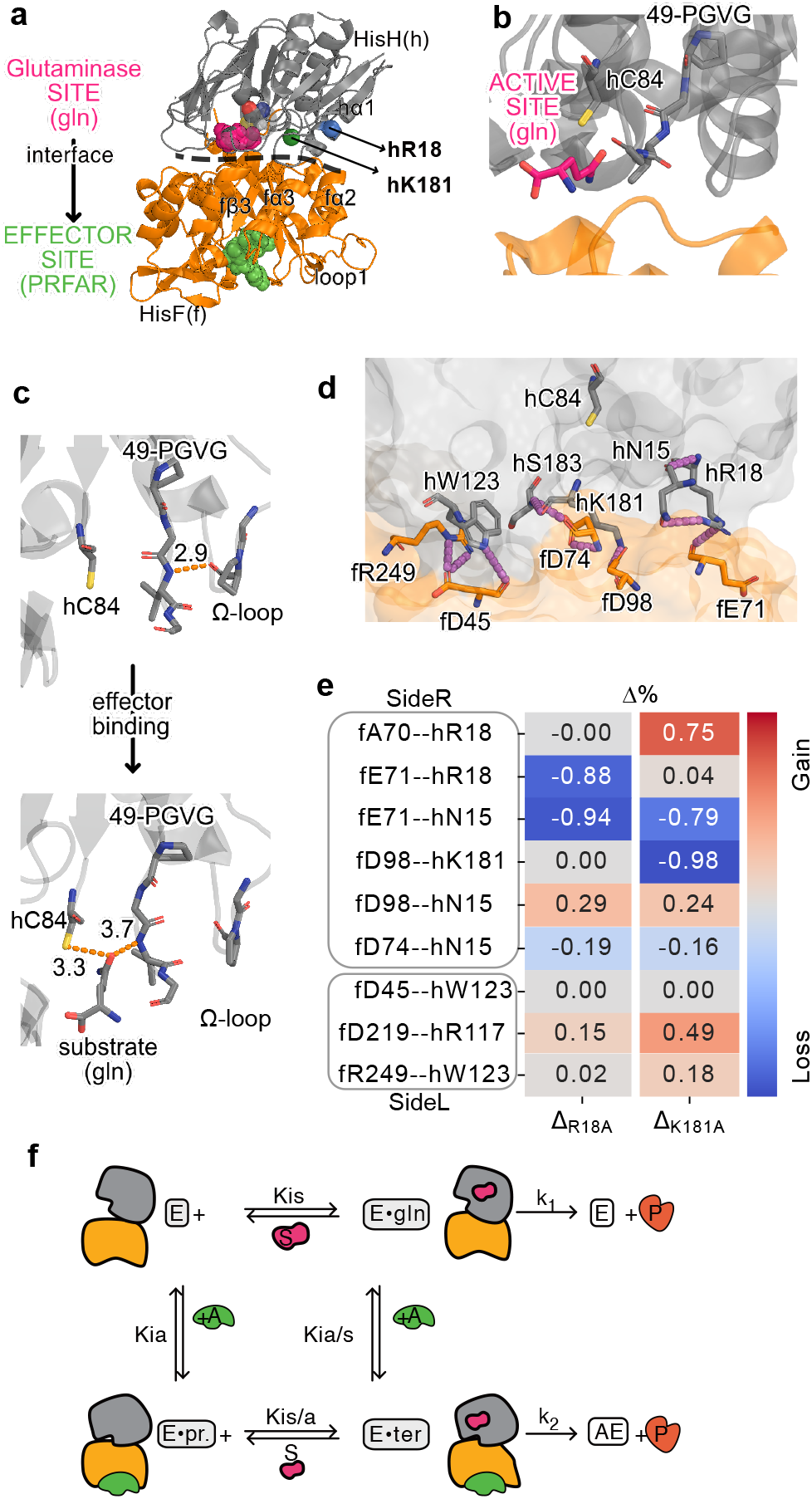
Structural basis of altered catalysis in IGPS. (a) Overall architecture of IGPS showing HisF, the cyclase domain, and HisH, the glutaminase domain. The interfacial region, residues mutated in this study (hR18 and hK181), and the catalytic cysteine (hC84) are shown. (b) Location of the glutaminase active site and the 49-PGVG sequence. (c) Rupture of the P10-V51 hydrogen-bond promotes formation of the catalytically competent state. (d) Interfacial noncovalent interactions in WT IGPS, including hydrogen bonds, salt bridges, and cation–*π* contacts between HisH residues (red sticks) and their HisF partners (green sticks). Red dashed lines indicate key contacts, including those involving hR18 and hK181. (e) Difference in interfacial contact probabilities for hR18A and hK181A relative to WT, averaged over concatenated MD trajectories across the four functional states and three replicates. Negative values indicate loss of contacts relative to WT (blue), positive values indicate gains (red). (f) Allosteric thermodynamic cycle of IGPS showing the four functional states analyzed by MD: apo enzyme (E), substrate-bound (E·gln), effector-bound enzyme (E·pr.), and ternary complex (E·ter). Equilibrium constants are defined as follows: Kis and Kia describe substrate (s = Gln) and effector (a) binding in the absence of the other ligand, respectively; Kis/a and Kia/s describe binding in the presence of saturating effector or substrate, respectively. IGPS is a V-type enzyme in which effector binding accelerates turnover of the E·gln complex, increasing k_2_ relative to k_1_.

The HisF–HisH interface contains a conserved network of hydrogen bonds, salt bridges, and hydrophobic contacts that transmit effector-induced conformational changes to the glutaminase catalytic site (Fig. 1d).(6)

Structural, spectroscopic, and computational studies have shown that effector binding stabilizes a closed interdomain interface, enhances *µ*s–ms dynamics across both subunits, and promotes catalytic turnover.(5, 13, 14) This interface forms a densely packed communication network that links the effector-binding site in HisF to the catalytic machinery in HisH. A charged interfacial gate formed by fR5, fE46, fK99, and fE167 regulates passage of ammonia generated by glutamine hydrolysis across the interface; perturbating this gate alters glutaminase activity.(5, 6, 13, 15) Residues near the interface, particularly within the f94–f99 region, move concertedly with the effector site during the allosteric transition(5, 16), suggesting that the interface serves as a mechanical and energetic hub that couples effector binding to catalysis.

Consistent with this model, perturbations at the interface can strongly influence catalytic activation. For example, the double mutant hK181A–fD98A removes a key salt-bridge interaction and disrupts the hinge motion that coordinates opening and closing of the interface, greatly reducing effector-induced activation.(17) The hK181A single substitution increases basal glutaminase activity,(18) whereas introduction of a light-switchable residue at the hinge position hW123 alters interface closure and enhances catalytic efficiency.(19) Together, these observations indicate that specific interfacial interactions act as control points that shape the conformational ensemble underlying allosteric regulation. Here, we investigate how targeted perturbations at the HisF–HisH interface switch the nature of the allosteric response. We focused on two single-site substitutions, hK181A and hR18A (Fig. 1a), which disrupt distinct salt-bridge networks linking HisF and HisH. By combining steady-state kinetics, solution NMR spectroscopy, and multi-*µ*s molecular dynamics simulations, we define the mechanistic basis for their divergent functional outcomes. hK181A increases basal activity and converts IGPS from a predominantly V-type allosteric enzyme into a mixed V+K-type enzyme, whereas hR18A disrupts communication across the interface and weakens effector-induced activation. These results show that specific interfacial residues act as molecular gates or communication hubs that determine how allosteric signals are transmitted, providing a strategy for rationally reprogramming enzyme regulation.

## Results

### Interface mutations produce divergent catalytic phenotypes and allosteric responses

To determine how targeted perturbations at the HisF–HisH interface can reprogram allosteric output, we examined two single-site substitutions, hK181A and hR18A, that disrupt distinct interfacial salt-bridge networks. We combined steady-state kinetics with molecular dynamics simulations of the four functional states of the IGPS cycle: apo enzyme (E), substrate-bound (E ·gln), effector-bound (E ·pr), and ternary complex (E ·ter) (Fig. 1f). Because WT IGPS behaves predominantly as a V-type allosteric enzyme, in which effector binding primarily enhances catalytic turnover, interface substitutions could either weaken this response or redistribute it toward a stronger K-type component. The two mutations produced distinct catalytic phenotypes (Fig. 2a,b). hK181A increased basal activity and, upon effector binding, showed a pronounced change in apparent substrate affinity together with the altered turnover, yielding a mixed V+K-type response. By contrast, hR18A impaired allosteric activation and caused only modest changes in catalytic efficiency relative to WT. Consistent with recent studies, including our own,(9, 19, 20) these findings indicate that the subunit interface acts as a regulatory layer that controls both the magnitude and mechanistic character of allosteric regulation in IGPS.

**Fig. 2.**
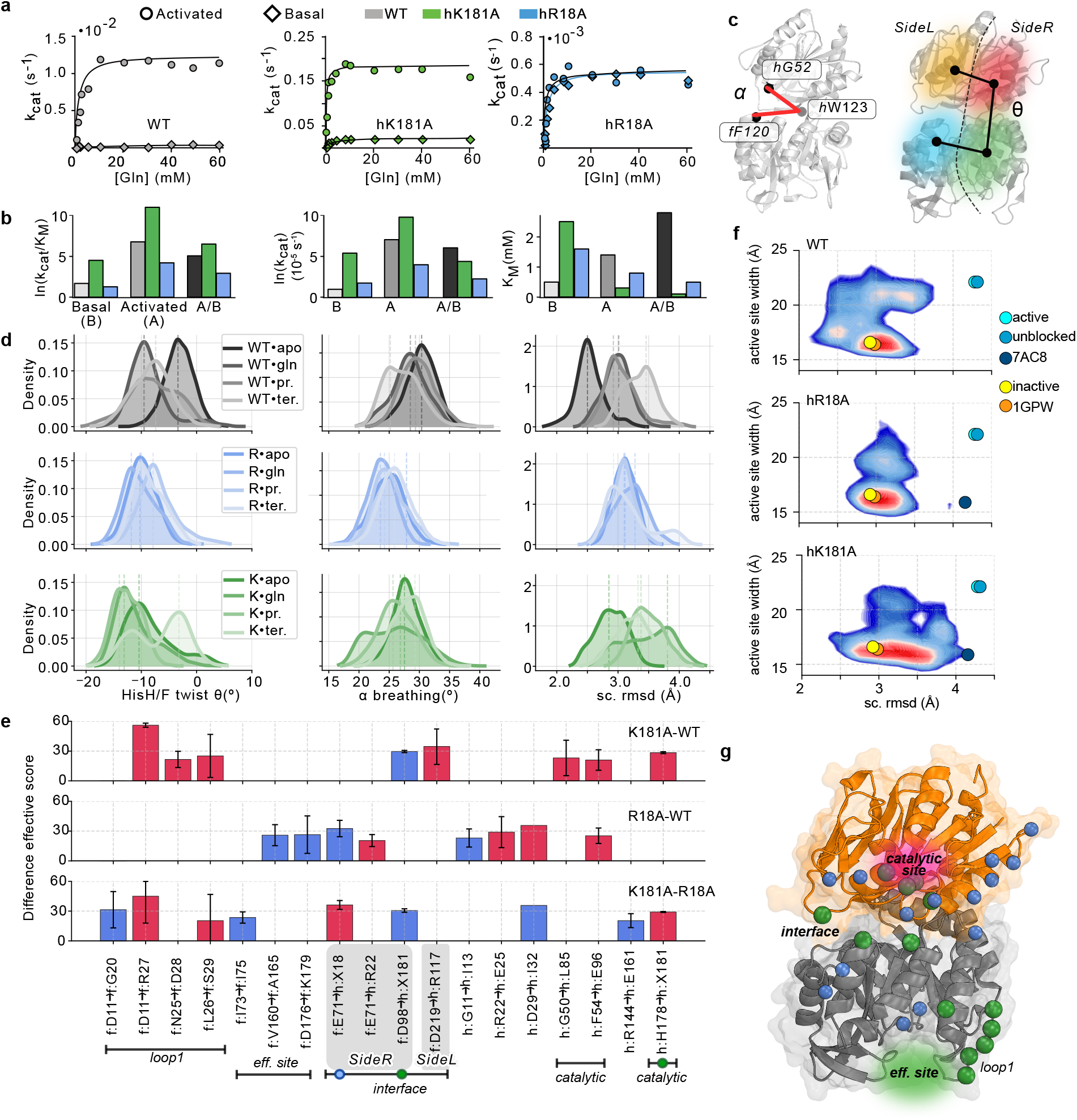
Kinetic response to interfacial mutation and associated interface dynamics. (a) Michaelis–Menten curves for WT, hR18A, and hK181A under apo and effector-bound conditions. Apo data are shown as diamonds and effector-bound data as circles. Unless otherwise indicated, WT is shown in gray, hK181A in green, and hR18A in blue. (b) Fold-change in catalytic efficiency (*k*_*cat*_*/K*_*m*_) upon PRFAR binding for WT and mutants. WT displays a predominantly V-type enhancement, hK181A shows a mixed V+K-type response, and hR18A exhibits impaired activation. (c) Collective variables used to monitor interface dynamics: the *α* hinge breathing angle and the HisF/HisH twist angle *θ*, defined as the dihedral formed by the centers of mass of the SideR and SideL subdomains. (d) Distribution of interface opening angles from MD trajectories, showing that both mutants preferentially sample closed states relative to WT, together with side-chain RMSD distributions highlighting differences in local *µ*s-timescale mobility across WT and mutant enzymes. (e) Largest changes in residue-pair effective noncovalent interaction scores at the HisH–HisF interface, active site, loop 1, and allosteric-activator-binding site (eff.site), computed for each mutant relative to WT. Blue bars indicate decreases interactions and red bars indicate increased interctions. Blue and green circles mark changes at the mutated sites. Details are provided in the Supplementary Information - *Non-covalent interaction analysis and delta effective scores*). (f) Conformational sampling of catalytic geometries in WT and mutant enzymes, shown as pseudo-free-energy histograms or density plots derived from MD simulations. Productive states are defined by alignment of the oxyanion-hole and active-site loops relative to a global RMS metric across apo, substrate-bound, effector-bound, and ternary ensembles. (g) Structural mapping of the largest hydrogen bond changes at the HisH–HisF interface, active site, loop1 and and PRFAR binding site.

### Structural context and interaction networks

HisF and HisH meet at a densely packed polar interface that acts as a conformational gate coupling effector binding in HisF to catalytic activation in HisH (Fig. 1a,d). In WT IGPS, hK181 forms a salt bridge with fD98 that is most populated in closed-interface conformations and is embedded in an electrostatic network that includes fD74 and hS183. This region is positioned to couple interfacial rearrangements to motions associated with oxyanion-hole formation in the HisH active site.(5, 12, 21) By contrast, hR18 forms a charge-pair interaction with fE71 near the interdomain hinge, within a polar network associated with hinge flexibility and intersubunit communication. Alanine substitutions at these sites perturb the interface in distinct ways. hK181A abolishes the hK181–fD98 salt bridge and weakens neighboring contacts within the same cluster, whereas hR18A removes the hR18–fE71 interaction and alters hinge-proximal hydrogen-bonding patterns. To quantify these effects, we computed changes in interresidue contact probabilities relative to WT across concatenated trajectories spanning the four functional states shown in Fig. 1f. hK181A primarily altered contacts centered on the fD98 region, together with compensatory gains involving nearby loop 1–HisF interactions, consistent with local reorganization of an otherwise coherent interface. In contrast, hR18A caused broad contact losses around the hinge region and more dispersed gains in distal HisF interactions, consistent with a less focused and more weakly coordinated communication network (Fig. 1e). Distance distributions for selected interfacial contacts and state-resolved contact persistence are provided in Fig. S1 and S2.

### Kinetic response to mutation and interface dynamics

Steady-state kinetics showed that hK181A increases basal catalytic turnover by nearly two orders of magnitude relative to WT (*k*_*cat*_ = 230 *×*10^*−*5^ s^*−*1^ vs. 2.7 *×*10^*−*5^ s^*−*1^), with a fivefold higher *K*_*m*_ in the apo enzyme, indicating reduced apparent substrate affinity (Fig. 2a). Upon activation by imidazole glycerol phosphate (IGP), hK181A showed a 680-fold increase in catalytic efficiency (*k*_*cat*_*/K*_*m*_), exceeding the 165-fold increase observed for WT (Fig. 2b). This enhanced response arose primarily from a pronounced decrease in *K*_*m*_ upon activation, indicating a strong K-type component superimposed on a V-type increase in *k*_*cat*_. Thus, hK181A redistributes the allosteric response from the predominantly V-type behavior of WT toward a mixed V+K-type profile.

By contrast, hR18A produces only modest changes in basal catalytic activity and markedly reduced activation. Upon effector binding, hR18A showed an approximately 10-fold increase in *k*_cat_ and a 20-fold increase in catalytic efficiency, defined as (*k*_cat_*/K*_*m*_)_act_*/*(*k*_cat_*/K*_*m*_)_basal_, substantially smaller than the activation observed for WT and hK181A, consistent with impaired allosteric coupling. Full kinetic parameters are reported in Table S1. In figures throughout the manuscript, WT is shown in gray, hK181A in green, and hR18A in blue.

### Interface configuration and hydrogen-bonding

To connect the kinetic phenotypes with structural dynamics, we analyzed the HisF–HisH interface using two collective variables previously shown to report on interdomain motion(13, 14): a hinge breathing angle (*α*) and an interdomain twist angle (*θ*). These coordinates are shown schematically in Fig. 2c.

Multi-microsecond MD simulations showed that both mutants preferentially sampled more closed apo-state interfaces than WT (Fig. 2d). However, the structural consequences of closure differed between variants. In hK181A, closed conformations were associated with stabilized contacts between loop 1 and the HisF barrel while preserving a coherent interfacial hydrogen-bond network. This behavior was accompanied by enrichment of catalytically productive geometries along the HisH–HisF twist coordinate *θ* (Fig. 2f).

By contrast, hR18A redistributes strain toward the hinge region. This perturbation altered the interfacial non-covalent interaction network, as reflected by changes in residue-pair effective interaction scores. hR18A also sampled a narrower range of twist angles, consistent with a stiffer dynamical profile that was largely insensitive to substrate or activation binding. Consequently, hR18A populated fewer productive catalytic configurations, even in the presence of the allosteric activator (Fig. 2e,f). In contrast, hK181A retained opening—closing fluctuations comparable to WT across multi-microsecond trajectories, consistent with its elevated basal *k*_*cat*_.

The altered flexibility of hR18A was further supported by side-chain mobility distributions, which showed reduced microsecond-timescale motion relative to WT and hK181A (Fig. 2d). Conversely, apo hK181A sampled closed-interface conformations, defined by smalled *α* values, more frequently than apo WT, consistent with its enhanced basal catalytic activity. These trends were robust to alternative definitions of the collective variables or atomic fluctuation metrics (SI, Fig. S3).

### NMR chemical shift perturbations reveal localized gates and distributed hubs

The kinetic and simulation results above suggest that catalytic efficiency depends not simply on interface closure, but also on how structural rearrangements redistribute communication pathways across the HisF–HisH interface. To experimentally localize these responses, we analyzed residue-resolved NMR chemical shift perturbations (CSPs) in the HisF subunit (Fig. 3).

**Fig. 3.**
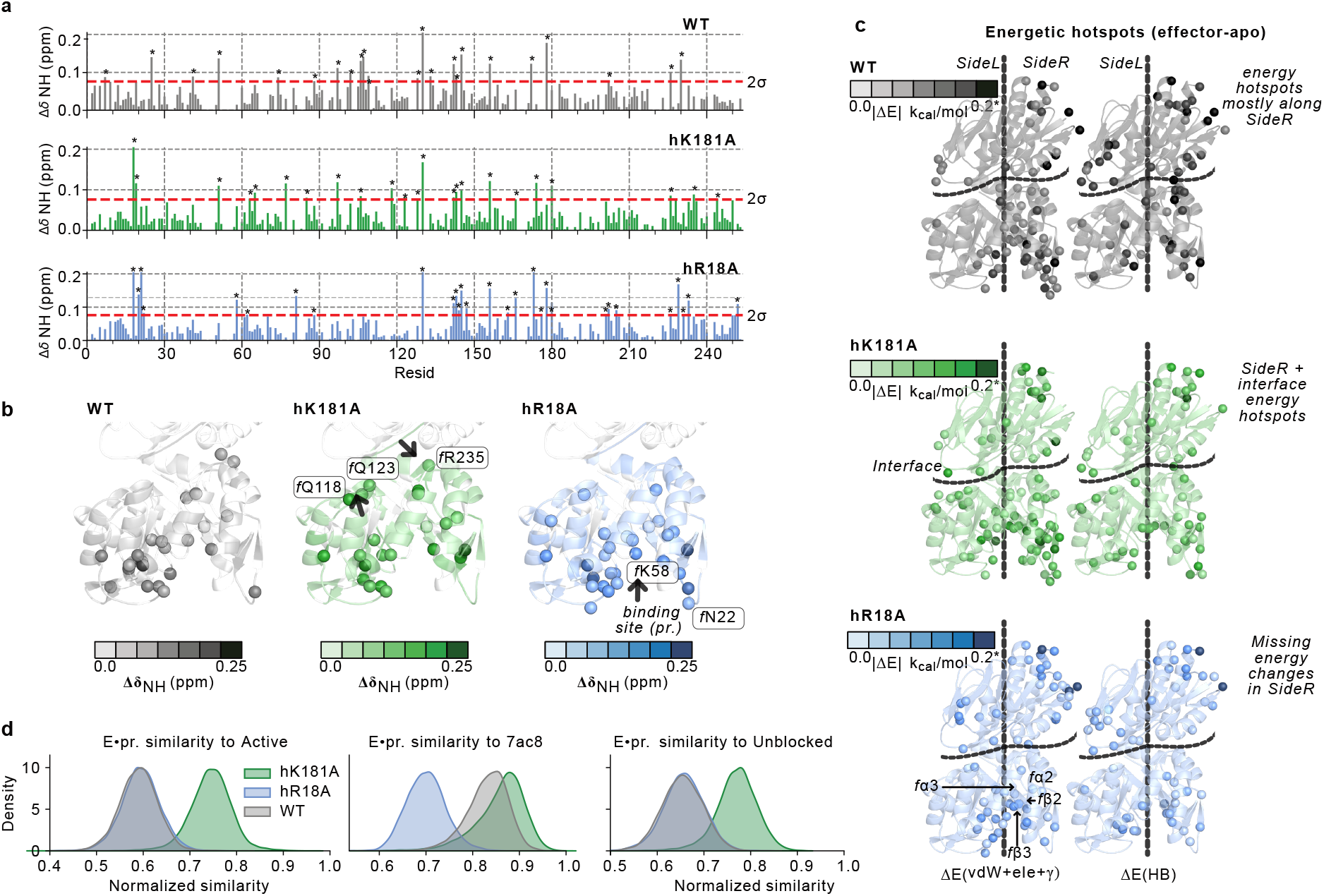
Site-resolved chemical-shift perturbations and energetic hotspots distinguish local “gates” from distributed “hubs”. (a) Backbone amide chemical-shift perturbations (CSPs) for WT·(effector–apo), hK181A·(effector–apo), and hR18A·(effector–apo), shown as residue-resolved traces. The red horizontal line indicates the threshold for significant perturbations, defined as the mean plus two standard deviations of the 10% trimmed CSP distribution. (b) CSP outliers indicated by asterisks are mapped onto the HisF structure. Outlier residues are colored according to perturbation intensity using mutant-specific palettes: gray for WT, green for hK181A, and blue for hR18A. (c) Energetic hotspots from multi-*µ*s molecular dynamics simulations, shown as effector-induced differences relative to the apo state, Δ*E* = *E*_pr_ −*E*_apo_. The left column reports the total per-residue energy change, Δ*E*_tot_, including electrostatic, van der Waals, nonpolar solvent-accessible surface area, and polar correction terms, displayed on a 0–2 kcal mol^*−*1^ scale. The total interaction energy (Δ*E*_tot_), is detailed in the Methods section. The right column shows the hydrogen-bond contribution, Δ*E*_hb_, displayed on a 0–0.2 kcal mol^*−*1^ scale. Values exceeding the upper limit of each scale were capped to improve visualization. (d) Distribution of conformational similarity between MD trajectories and reference structural states. Reference conformations correspond to the active state and an “unblocked” intermediate displaying a partially flipped oxyanion strand,(21) as well as the active crystal structure (PDB ID: 7AC8).(14) Similarity was computed from Mahalanobis distances in the space of structural metrics describing interface geometry, active-site configuration, and interdomain motions. Distances for each trajectory frame were normalized to a 0–1 similarity scale (1 = closest to the reference). Kernel density estimates show the distribution of similarity values for WT (gray), hK181A (green), and hR18A (blue) in effector-bound ensembles.

We compared ^1^H–^15^N HSQC spectra for WT, hK181A, and hR18A in the effector-bound state with their corresponding apo spectra. CSP values for WT · (effector–apo), hK181A ·(effector–apo), and hR18A · (effector–apo) are shown as residue-wise traces in Fig. 3a. Significant perturbations were defined as CSP outliers exceeding the 10% trimmed mean plus two SDs of the CSP distribution (see Methods) and mapped onto the HisF structure with color intensity proportional to CSP magnitude (Fig. 3b).

Three features emerged. First, all variants showed pronounced CSPs near the effector-binding pocket, including HisF residues 173–180, consistent with direct ligand engagement. Second, WT and hK181A showed additional perturbations at the HisF–HisH interface, including fR235. In hK181A, these perturbations extended to a reinforced cluster spanning fQ118–fQ123. These localized responses coincide with structural elements implicated in productive interface closure and are consistent with the enhanced apparent substrate affinity and mixed *V* and *K*-type activation observed for hK181A. Third, hR18A showed fewer and weaker CSPs, with perturbations largely confined to the activator-binding pocket and limited propagation toward the interdomain contact region.

This pattern is consistent with impaired transmission of the allosteric signal from the activator site to the catalytic subunit, explaining the attenuated activation observed kinetically.

Together, these data support distinct mechanisms for the two mutants. hK181A enhances catalysis by reinforcing a localized regulatory gate at the interface, whereas hR18A weakens a distributed communication hub required for efficient allosteric signal propagation.

### Energetic hotspots explain clustered CSPs in hK181A and dispersed CSPs in hR18A

To relate the CSP patterns to interaction energetics, we computed per-residue interaction energies from multi-microsecond MD trajectories for each state and variant. Effector-induced changes were evaluated relative to the apo ensemble as Δ*E* = *E*_effector_ *− E*_apo_.

We analyzed two complementary energetic metrics. The first, Δ*E*_tot_, reports the total per-residue interaction score, including electrostatics, van der Waals interactions, a nonpolar SASA term (*γ*_SASA_), and a continuum electrostatic correction. The second, Δ*E*_hb_, reports directional hydrogen-bond contributions, with individual contacts capped at 0.8 kcal mol^*−*1^ to avoid overweighting transient interactions. Full details are provided in Supplementary Information - **Energy Decomposition Analysis**). Hotspots in Δ*E*_tot_ and Δ*E*_hb_ are shown in Fig. 3c.

Using the established SideL/SideR partition of HisF dynamics (5, 22), WT IGPS showed effector-induced energetic hotspots extending from the effector-binding pocket across SideR, particularly through the *β*2–*β*3 and *α*2–*α*3 elements and loop 1—toward the HisH interface (Fig. 1a). This corridor corresponds to the canonical SideR communication pathway implicated in catalytic activation.(5)

hK181A preserved this energetic corridor but introduced a focused redistribution of interactions at the interface. Energetic changes were concentrated near the disrupted hK181—fD98 salt-bridge region and neighboring contacts, including fQ123 and the fR191–PLTT–fL196 segment, with HisH residues such as hR367 and hR370. Notably, hR18 emerged as a significant energetic hotspot in both WT and hK181A for Δ*E*_tot_ and Δ*E*_hb_. In hR18A, this hotspot was largely lost, with only a residual hydrogen-bond contribution remaining, consistent with the disruption of the interfacial salt bridge and reduced coupling.

By contrast, hR18A showed a markedly different energetic pattern. Rather than forming a continuous SideR corridor from the effector pocket to the HisF–HisH interface, hotspots shifted toward SideL elements surrounding the binding pocket and showed limited propagation toward HisH. This fragmented pattern is consistent with weakened long-range communication across the interface.

### CSP energy convergence identifies gates and hubs that tune catalytic output

Comparison of CSP outliers with energetic hotspots further distinguished the two mutants. In hK181A, residues showing both strong CSP and energetic perturbations clustered at the HisF–HisH interface, forming a contiguous patch that involved loop 1 and the *β*8–*α*4 region. This interface-centered cluster behaves as a localized regulatory gate whose reweighting stabilizes productive closed conformations.

In hR18A, residues showing combined CSP and energetic perturbations are instead scattered across multiple structural elements near the effector pocket and hinge region. Rather than converging at the interface, these perturbations define dispersed communication nodes, or distributed hubs, whose weakening disrupts coherent signal transmission.

These patterns provide a mechanistic basis for the distinct kinetic responses of the two variants. Perturbations localized at the interfacial gate appear correlated with changes in substrate affinity. hK181A, which reinforced this cohesive interface region, showed the largest activation-dependent decrease in *K*_*m*_. In contrast, hR18A underwent geometric interface closure but lacked cohesive energetic coupling at the gate, and therefore showed little change in substrate affinity or catalytic output.

Similarly, energetic continuity along the SideR corridor appears to be associated with catalytic turnover. WT and hK181A maintain this mixed hydrophobic and electrostatic communication pathway communication pathway and display stronger *k*_*cat*_ responses, whereas hR18A disrupts the corridor and exhibits reduced catalytic activation.

This mechanistic interpretation is further supported by ensemble-level analysis of trajectory similarity to different reference states (i.e, a crystal structure of the activated state(14), a modeled active-state and partially activated one(21), Fig. 3d). Mahalanobis-distance similarity distributions show that hK181A samples conformations closer to active-like reference structures than WT, whereas hR18A remains shifted toward less productive regions of conformational space. Together, these results indicate that interfacial gate cohesion primarily tunes substrate affinity (*K*_*m*_), while long-range SideR communication governs catalytic turnover (*k*_*cat*_), allowing targeted interface mutations to redistribute the balance between V-type and K-type allosteric responses. Consistent with this interpretation, analysis of catalytic-site geometry in the simulations indicates that hK181A samples conformations compatible with formation of the oxyanion hole (involving hG50) more frequently than WT, whereas such configurations remain comparatively absent in hR18A (Fig. S4).

### Altered *µ*s–ms dynamics and communication networks explain mutation-specific allosteric responses

We next examined whether the structural and energetic perturbations identified above translate into changes in *µ*s–ms conformational dynamics, the timescale most closely associated with allosteric activation in IGPS. Methyl ^13^C relaxation-dispersion experiments were used to quantify chemical exchange broadening across the protein in the absence and presence of the allosteric effector IGP (Fig. 4a,b).

**Fig. 4.**
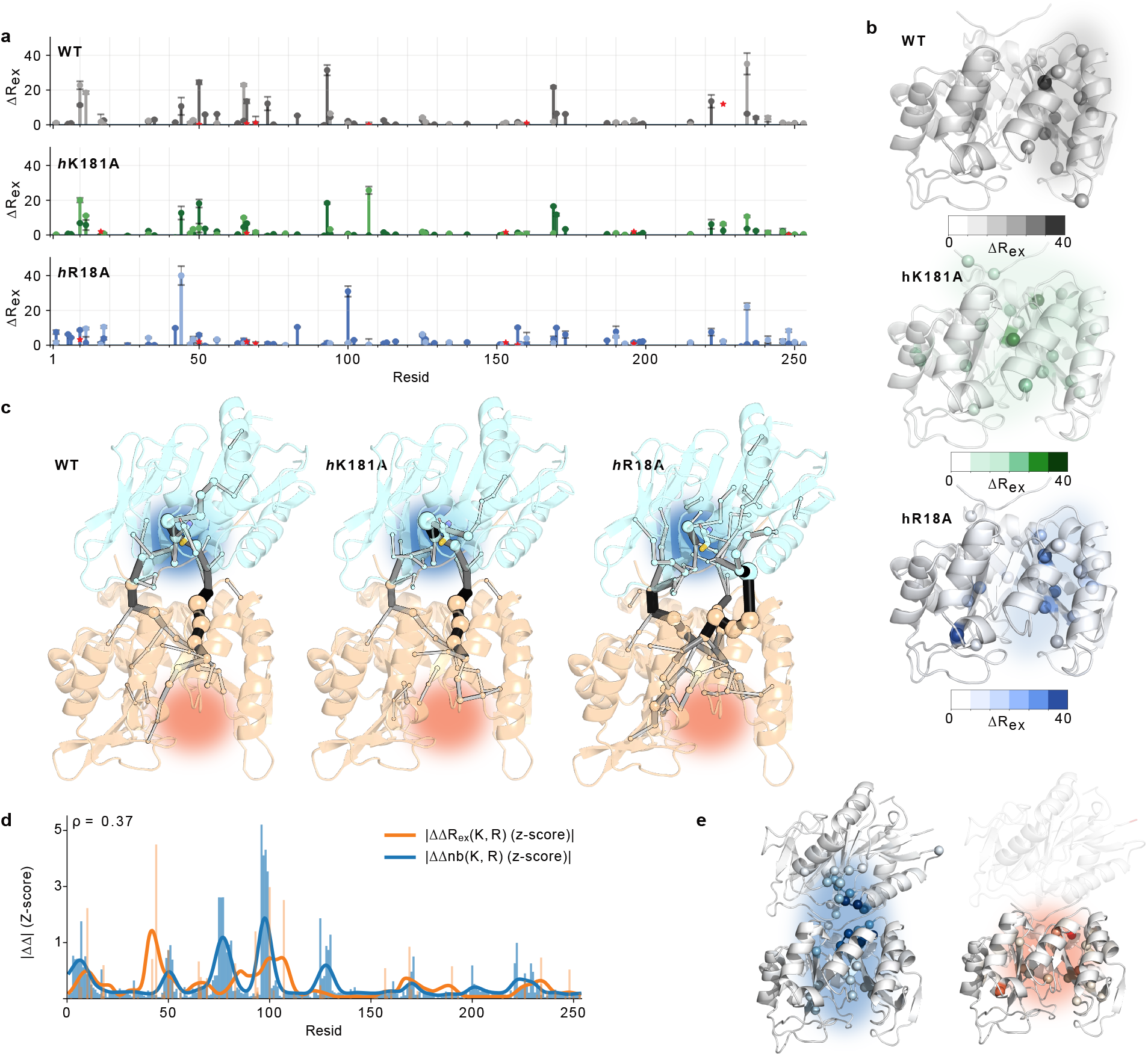
Distinct responses to effector binding in WT and interface mutants. Effector binding alters the dynamics and structural ensembles of IGPS in variant-specific ways. (a) Residue-resolved estimate of exchange broadening extracted from methyl CPMG relaxation–dispersion profiles. For each probe, the exchange-sensitive amplitude was computed as Δ*R*_ex_ = (⟨*R*_2_⟩_low_ − ⟨*R*_2_⟩_high_)_effector_ − (⟨*R*_2_⟩_low_ − ⟨*R*_2_⟩_high_)_apo_, where ⟨*R*_2_⟩_low_ and ⟨*R*_2_⟩_high_ are the mean transverse relaxation rates measured at the lowest and highest CPMG pulsing frequencies, respectively. The averages were evaluated over the *k* lowest and *k* highest CPMG pulsing frequency points of each dispersion profile. Uncertainties were propagated from the reported standard deviations by first computing the error of each mean, 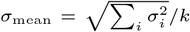, and then combining the low- and high-frequency contributions in quadrature. Values are reported separately for each residue and proton type. Negative values, corresponding to an absence of detectable *µ*s–ms exchange broadening under these conditions, are marked by red stars. (b) Structural mapping of residue-resolved methyl relaxation-dispersion amplitudes onto the IGPS structure. For each condition, the quantity plotted on the protein is the residue-specific Δ*R*_ex_ *>* 0 value obtained from the corresponding holo–apo comparison and assigned to the C*α* atom of each residue. Cartoon representations are colored using condition-specific sequential color scales, with larger positive Δ*R*_ex_ values shown in darker shades and residues without assigned values left at the baseline color. Spheres indicate the top 20 residues with the highest positive Δ*R*_ex_ values in each dataset. No explicit numerical cutoff was applied beyond restricting sphere display to positive values only; all available residue values were used for the coloring, whereas only the 20 largest positive values were highlighted as spheres. (c) Allosteric communication pathways inferred from dynamical correlation networks constructed from concatenated 12-*µ*s molecular dynamics trajectories (3 *µ*s per state: E, E·pr, E·gln, E·gln·pr) for WT, hK181A, and hR18A. Residues are represented as nodes connected by edges weighted by generalized linear mutual information (GLMI) between backbone dihedral fluctuations. Shortest-path edge betweenness centrality was computed on the resulting weighted networks to identify the most frequently traversed routes of signal transmission. Edge thickness reflects normalized edge betweenness, while node size reflects normalized node betweenness. For visualization clarity, only edges with normalized betweenness greater than 0.2 are shown. The resulting traffic maps highlight dominant allosteric communication routes within the protein: hK181A largely preserves or strengthens communication across the interfacial and SideR regions, whereas hR18A displays weaker and more fragmented pathways relative to WT. (d) Residue-wise comparison between experimental and simulation-derived differential allosteric responses. The bar plots show ΔΔ*R*_ex_(*K, R*) together with ΔΔ*n*_*b*_(*K, R*), corresponding to the difference between hK181A and hR18A in experimental methyl relaxation-dispersion amplitudes and in network node betweenness, respectively. Continuous curves represent Gaussian-broadened versions of the residue-wise profiles, used to emphasize sequence-localized trends. For the ΔΔ*n*_*b*_(*K, R*) comparison, both profiles were min–max scaled before smoothing; the ΔΔ*e*_*b*_(*K, R*) comparison is shown without scaling. The Spearman coefficient *ρ* quantifies the correlation between the two curves. (e) Structural representations show ΔΔ*n*_*b*_(*K, R*) (left) and ΔΔ*R*_ex_(*K, R*) (right) mapped onto the IGPS structure, highlighting regions where experimental differential exchange and differential network traffic changes colocalize.

In WT IGPS, effector binding increases the amplitude of exchange broadening for a cluster of residues spanning the HisF *α*2*/α*3 helices and the surrounding hydrophobic core (Fig. 4a). Mapping these residues onto the structure reveals a contiguous region of enhanced dynamics connecting the effector pocket to the interdomain interface (Fig. 4b). This pattern is consistent with previous work linking *µ*s–ms motions in these elements to productive interface closure and catalytic activation.(13, 16, 23)

The hK181A mutation preserves and partially amplifies this dynamic response. Several residues that show conformational exchange only in the effector-bound WT enzyme already display measurable exchange in the apo hK181A variant, indicating that the mutation pre-organizes the enzyme toward catalytically competent conformations. Effector binding further increases the magnitude of exchange contributions in this region, producing a dynamic network closely resembling that observed in WT but shifted toward greater baseline flexibility. This redistribution of conformational sampling is consistent with the enhanced basal activity and reduced *K*_*m*_ observed for hK181A.

By contrast, hR18A exhibits markedly weaker exchange responses. Only a small number of residues display measurable Δ*R*_ex_ changes upon effector binding, and these residues do not form the contiguous hydrophobic-core cluster observed in WT and hK181A (Fig. 4a,b). Instead, dynamic perturbations remain sparse and localized near the effector pocket. This muted response mirrors the diffuse CSP profile and fragmented energetic hotspots identified in Fig. 3, indicating that the mutation disrupts the propagation of dynamic signals from the effector-binding site to the catalytic interface.

To understand how these dynamic differences translate into long-range communication, we constructed residue-level dynamical correlation networks from the concatenated MD trajectories for each mutant. Edge weights were derived from generalized mutual information between backbone dihedral fluctuations, and node and edge betweenness centralities were used to identify dominant communication routes. The resulting traffic maps were projected onto the protein structure (Fig. 4c), following a procedure similar to that described in Ref. (24, 25).

The hK181A mutation preserves the overall architecture of this communication network while strengthening traffic through the hinge-mediated branch. Enhanced betweenness along *fα*3, *fβ*3 and adjacent structural elements indicates that this pathway partially compensates for the disrupted hK181–fD98 contact, maintaining efficient coupling between the effector and catalytic sites. In contrast, hR18A substantially weakens both interfacial and hinge-mediated communication routes. Traffic becomes distributed across longer and less direct paths within the HisF barrel, resulting in a fragmented network with fewer high-betweenness edges connecting the effector and catalytic regions.

Direct comparison between experiment and simulation highlights this correspondence. Residue-wise differences in experimental exchange amplitudes between hK181A and hR18A (ΔΔ*R*_ex_) correlate with differences in network node betweenness derived from the MD simulations (ΔΔ*n*_*b*_) (Fig. 4d). Regions exhibiting stronger exchange in hK181A coincide with residues that carry increased communication traffic in the simulated networks. Structural mapping of these differential signals further demonstrates that experimental dynamic hotspots colocalize with the network pathways identified computationally (Fig. 4e).

Taken together, these results establish a consistent mechanistic picture across experimental and computational observables. Local structural perturbations at the HisF–HisH interface (Fig. 3) propagate into changes in *µ*s–ms dynamics (Fig. 4a,b) and ultimately reshape the global communication network linking effector binding to catalysis (Fig. 4c–e). The hK181A mutation preserves a cohesive communication corridor while strengthening hinge-mediated pathways, producing an enzyme with enhanced basal activity and a mixed V+K-type response. In contrast, the hR18A mutation disrupts key interfacial contacts that normally coordinate long-range signaling, yielding a fragmented communication network and attenuated catalytic activation.

These findings demonstrate that specific interfacial residues function as regulatory control points within the allosteric network of IGPS. By selectively perturbing these sites, it is possible to redistribute communication traffic through the protein and thereby tune the balance between V-type and K-type allosteric responses.

## Discussion

The divergent behavior of hK181A and hR18A shows that the HisF-HisH interface is not a uniform conduit for allosteric signaling, but a structured regulatory surface in which nearby contacts play distinct mechanistic roles. By integrating steady-state kinetics, NMR spectroscopy, and multi-*µ*s molecular dynamics simulations, we show that two substitutions in HisH that disrupt distinct salt-bridge interactions—hK181A and hR18A—produce opposite allosteric outcomes. The hK181A mutation converts the predominantly V-type activation of the wild-type enzyme into a mixed response with a strong K-type component, whereas hR18A attenuates activation altogether. These divergent phenotypes arise from distinct reorganizations of the allosteric communication network linking the effector-binding pocket in HisF to the catalytic site in HisH.

In hK181A, disruption of the hK181–fD98 salt bridge produces a localized energetic reorganization at the interface. CSP measurements identify perturbations in loop 1 and the *β*8–*α*4 region of HisF, while MD simulations reveal that the interface samples closed conformations even in the absence of effector. Despite the loss of the original salt bridge, communication through the interface is preserved through a reinforced hinge-mediated pathway, delocalized over the interface region. This alternative route maintains efficient coupling between the effector pocket and the catalytic region, consistent with the enhanced baseline *µ*s–ms dynamics and the preserved allosteric traffic observed in the simulations. Functionally, this redistribution is consistent with a conformational ensemble biased toward substrate-affine states, explaining the pronounced decrease in *K*_*m*_ upon activation while maintaining substantial catalytic turnover. In this sense, hK181 acts as a local regulatory gate whose perturbation reweights the ensemble without disrupting the overall communication network.

By contrast, the hR18A mutation eliminates the hR18–fE71 interaction near the interdomain hinge, producing diffuse energetic perturbations and broadly distributed CSPs that fail to converge at the interface. Relaxation-dispersion experiments reveal fewer residues undergoing *µ*s–ms exchange and a muted dynamic response to effector binding. Network analysis further shows that the loss of this hinge contact fragments the communication architecture, weakening both the direct interfacial pathway and the hinge-mediated route connecting the effector site to the catalytic loop. As a result, effector binding cannot efficiently repopulate catalytically competent conformations, leading to attenuated activation. This behavior highlights the distinct role of hR18 as a hub within the communication network: unlike the gate-like role of hK181, perturbation of this residue disrupts distributed pathways across the protein.

Taken together, these results reveal that the HisF–HisH interface functions as a tunable control point for IGPS allostery. Individual interfacial contacts can selectively bias how signals propagate through the communication network, thereby shifting the balance between V-type and K-type regulation. Mutations such as hK181A preserve network integrity while redistributing traffic through alternative pathways, whereas mutations such as hR18A disrupt key communication hubs and reduce signaling efficiency.

More broadly, this work illustrates the power of combining experimental observables with atomistic simulations to resolve the hierarchical layers of allosteric regulation—from local energetic perturbations to global communication networks. Our analysis shows how single-residue changes reshape the conformational ensemble and thereby alter kinetic outputs, even within an otherwise conserved structural scaffold. By linking structural perturbations, energetic hotspots, *µ*s–ms dynamics, and network traffic to measurable kinetic effects, we establish a multiscale framework for understanding how mutations reprogram enzyme regulation.

Finally, the principles identified here resonate with a broader shift in the study and engineering of allosteric systems. Rather than viewing allostery primarily as a structural transition between discrete states, recent work increasingly emphasizes the role of communication networks and energetic landscapes that govern conformational ensembles.(26) Network-based analyses have shown that specific residues can act as critical communication nodes whose perturbation alters long-range signaling and functional output.(27) Similarly, large-scale energetic profiling of PDZ domains demonstrated that structural extensions reshape the allosteric landscape by altering energetic couplings among distant residues.(28, 29). Computational and NMR analyses have likewise revealed conserved residue networks that propagate activation signals across large structural distances.(30)

In parallel, several studies have begun to exploit these principles to engineer allosteric behavior. For example, manipulation of conformational equilibria and communication pathways has enabled the tuning of catalytic regulation in metabolic enzymes such as phosphofructokinase,(31) while semi-rational redesign of the IGPS interface has been used to amplify light-controlled catalytic responses through engineered conformational shifts.(20) Together, these studies show that allosteric regulation can be modified by targeted perturbations, but they do not always reveal why a given perturbation succeeds, fails, or produces a specific regulatory outcome.

Our results provide this mechanistic link. We show that single interfacial residues can play distinct roles within the allosteric network: some act as local gates that stabilize productive conformational states, whereas others act as hubs that coordinate long-range communication across the enzyme. As a result, interface mutations do not simply weaken or strengthen allostery in a nonspecific manner. Instead, they redistribute communication traffic, energetic coupling, and *µ*s–ms dynamics in ways that determine whether the enzyme retains V-type regulation, gains a K-type component, or loses efficient activation. In this sense, the HisF-HisH interface of IGPS provides a concrete example of how network-level reasoning can be translated into rational control of enzyme regulation. By identifying the mechanistic role of individual interfacial residues, this work shows how interface reengineering can be used to rationally reprogram allosteric output rather than relying on serendipitous functional screening.

## Materials and Methods

### Molecular dynamics simulations

Molecular dynamics (MD) simulations were performed for the apo enzyme, substrate-bound (E•gln), effector-bound (E•pr), and ternary (E•ter) states of *Thermotoga maritima* IGPS starting from the crystal structure (PDB ID: 1GPW).(6) Mutant systems (hR18A and hK181A) were generated by single-site substitution in the HisH subunit. Systems were parameterized using the Amber ff19SB force field(32) together with GAFF parameters(33) for PRFAR, solvated in explicit water, neutralized with counterions, and simulated using the AMBER GPU implementation under periodic boundary conditions.(34, 35) For each mutant and functional state, three independent trajectories of approximately 1.5 *µ*s were generated. After equilibration, the final 1 *µ*s (∼10,000 frames) of each trajectory was used for analysis, yielding 12 *µ*s of analyzed simulation per mutant across the four functional states. Detailed system preparation, equilibration procedures, and simulation parameters are described in Supplementary Information - *Molecular dynamics simulations*.

### Traffic analysis

Dynamical communication within IGPS was analyzed using correlation networks derived from MD trajectories, using the MDiGest python tool.(36) In this representation, residues are treated as nodes located at each residue position and edges encode dynamical correlations between residue motions, quantified using generalized mutual information (MI). Edges were defined only between residue pairs with any heavy-atom distance *<* 6 Å; all other pairs were assigned a weight of zero. For each such pair, the edge weight was set to *−* log(MI) of the backbone dihedral fluctuations, transforming the correlations into effective distances, and used to construct weighted residue interaction networks.

Shortest-path–based metrics were then used to quantify communication traffic through the network. In particular, node and edge betweenness centrality were computed on the weighted graphs as implemented in NetworkX(37) to identify residues and residue pairs that participate most frequently in dominant communication pathways. Centrality values were normalized independently within each system and mapped onto the protein structure for visualization. A detailed description of network construction, weighting schemes, and visualization procedures is provided in the Supplementary Information.

### Non-covalent interaction analysis

Non-covalent interactions were computed from MD trajectories at the atom level for every analyzed frame and subsequently aggregated into residue pairs. Hydrogen bonds, hydrophobic contacts, and *π*–*π* stacking interactions were detected using geometric criteria, and their occupancies were evaluated over the trajectory.

For each residue pair, we defined an effective interaction score combining the mean occupancy of contributing contacts with their multiplicity. Scores were computed independently for each mutant, conformational state, and replicate, and averaged across the three simulation replicas. Differences between mutants (Δ scores) were then computed for each state and summarized across states to quantify mutation-induced perturbations of the interaction network.

Full details of interaction definitions, score construction, and statistical analysis are provided in the Supplementary Information - *Non-covalent interaction analysis and delta effective scores*.

### Expression and Purification of IGPS Mutants – Expression

Plasmids containing *E. coli*-optimized genes encoding HisH and HisF from *Thermotoga maritima*, cloned into pET43.1b vectors, were purchased from GenScript. Plasmids encoding the HisH sequence were used to generate desired mutants via PCR. The plasmids coding for the HisF sequence and the mutant HisH sequence (containing a C-terminal histidine tag) were co-transformed into BL21(DE3) cells (New England Biolabs)” since the histidine tag is actually on the HisH subunit. Isotopically enriched samples were prepared by overexpressing the HisF subunit in 1 L of deuterated M9 minimal medium containing ^15^NH_4_Cl (Cambridge Isotope Labs, MA) as the sole nitrogen source. The HisH subunit was overexpressed separately in 1.5 L of deuterated M9 minimal medium with naturally abundant nitrogen isotopes. Cell cultures were induced with 1 mM IPTG (isopropyl *β*-D-1-thiogalactopyranoside, Sigma-Aldrich) at OD_600_ of 0.8–1.0 and shaken at 30 ^*°*^C for 14–16 h. To incorporate ^13^C into the methyl groups of Ile, Leu, and Val (ILV) residues, 60 mg/L of *α*-ketobutyric acid, sodium salt (methyl-^13^C, 99%; 3,3-D_2_, 98%) and 100 mg/L of *α*-ketoisovaleric acid, sodium salt (3-methyl-^13^C, 99%; 3,4,4,4-D_4_, 98%) were added 30 min prior to induction.

### Purification

Cells expressing HisF or HisH were harvested at 4,000 rpm for 35 min, resuspended in lysis buffer (10 mM Tris-HCl, 10 mM CAPS, 300 mM NaCl, 1 mM *β*-mercaptoethanol, pH 7.5), and co-lysed by ultrasonication. The lysate was incubated at 333 K for 30 min and centrifuged to remove debris. The supernatant was filtered (0.22 *µ*m) and loaded onto Ni–NTA agarose resin equilibrated with lysis buffer. After 20 min incubation at 4 ^*°*^C, the column was washed with 12 column volumes of wash buffer (10 mM Tris–HCl, 10 mM CAPS, 300 mM NaCl, 1 mM *β*-mercaptoethanol, 15 mM imidazole, pH 9.5) and eluted with 8 column volumes of elution buffer of the same composition with 250mM imidazole. Fractions containing IGPS were pooled, dialyzed overnight against 2 L of NMR buffer (10 mM HEPES, 10 mM KCl, 0.5 mM EDTA, pH 7.3), concentrated, and supplemented with ∼8% D_2_O for NMR analysis.

### Steady-State Kinetic Studies

The glutaminase activity of wild-type and mutant IGPS was monitored as a function of glutamine concentration using a coupled assay adapted from Kneuttinger *et al*.(19) The glutaminase reaction produces NH_3_ and glutamate, which are converted by glutamate oxidase (GOX, 99%, Sigma-Aldrich) to *α*-ketoglutarate and H_2_O_2_. This product is coupled to horseradish peroxidase (HRP, 99%, Sigma-Aldrich), which, with phenol and 4-aminoantipyridine (4-AAP), forms a red quinoneimine chromophore with *ε*_505_ = 6,400 M^*−*1^cm^*−*1^. Reactions were conducted in 20 mM Tris–HCl (pH 7.0) containing 0.15 g/L HRP, 20 mU/mL GOX, 1 mM 4-AAP, 1 mM phenol, and 35–50 *µ*M enzyme for basal activity. Glutamine concentration ranged from 0.2–40 mM. Activated rates were measured in the presence of IGP (D-erythro-imidazoleglycerol phosphate monohydrate, 90%, Toronto Research Chemicals) using 200 mU/mL GOX and 3–5 *µ*M enzyme under otherwise identical conditions. Reactions were monitored at 505 nm using a BioTek Synergy 2.0 plate reader for 3 h at 30 ^*°*^C. Initial velocities were fitted to the Michaelis–Menten equation, and kinetic parameters were analyzed using Microsoft Excel and GraphPad Prism 9.(38)

### Solution NMR Studies – 2D ^1^H −^15^N TROSY-HSQC

All spectra were acquired at 600 MHz (^1^H) on a Varian Inova spectrometer at 30 ^*°*^C, using 32 scans and 128 increments in *t*_1_. Spectral widths were 12,000 Hz (direct) and 2,500 Hz (indirect), with a 1 s recycle delay. Carrier frequencies were set to water (^1^H) and 120 ppm (^15^N). WT chemical shifts (16) served as reference for mutant assignments. Spectra were processed with NMRPipe(39) and analyzed with Sparky.(40)

### ^13^CH_3_ Methyl-TROSY CPMG

Multiple-quantum CPMG dispersion experiments probing relaxation rates of ILV methyl groups were performed at 600 MHz on a Varian Inova spectrometer using the pulse sequence of Korzhnev *et al*.(41) The ^13^C carrier was set to 19.5 ppm (3,200 Hz width), and ^1^H carrier to 0.75 ppm (8,503 Hz width). Forty scans and 128 *t*_1_ increments were collected. The total relaxation delay was 40 ms with *τ*_cp_ points of 0.0, 0.40, 0.50, 0.625, 0.833, 1.0, 1.25, 1.667, 2.0, 2.5, 5.0, and 10 ms, and a 2 s recycle delay. Data were processed in NMRPipe(39) and analyzed in Sparky(40) with in-house scripts for relaxation-rate extraction.

### Chemical Shift Perturbation (CSP) Analysis

CSPs were computed as:

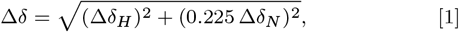

where Δ*δ*_*H*_ and Δ*δ*_*N*_ are the changes in ^1^H and ^15^N chemical shifts, respectively. Residues with Δ*δ* exceeding the 10% trimmed mean plus two standard deviations were classified as significantly perturbed. Additional details on experimental setup are provided as Supplementary Information.

### Energy Decomposition Analysis

Per-residue interaction energy decomposition was performed on dried MD trajectories (protein and ligands only) to localize energetic changes associated with mutation and effector binding. Each saved frame was analyzed using MDAnalysis(42) for atom selections and ParmEd(43) for topology and force-field parameters. Pairwise atom interactions were computed and subsequently aggregated at the residue level. **Electrostatic interactions** were evaluated using a distance-dependent dielectric model,

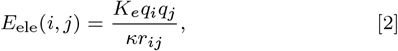

where *K*_*e*_ = 332.0636 kcal Å mol^*−*1^ e^*−*2^ and *κ* = 5.0.

**Van der Waals** interactions were computed using a Lennard– Jones potential with a smooth switching function applied between *r*_*s*_ = 8.0 Å and *r*_*c*_ = 9.5 Å.

**Hydrogen bonds** were modeled using an orientation-dependent term

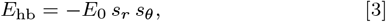

with *E*_0_ = 0.8 kcal*/*mol and geometric criteria H … A ≤ 2.5 Å and donor–H–acceptor angle ≥140^*°*^.

**Implicit solvent corrections** included a screened Coulomb term

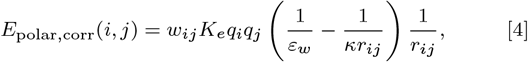

with *ε*_*w*_ = 78.5, and a nonpolar contribution proportional to the solvent-accessible surface area,

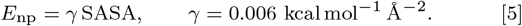

**Total interaction energy** was defined as

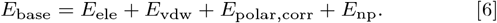

**Mutation effects** were quantified as

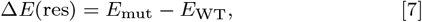

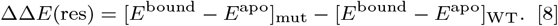

Unless otherwise noted, parameters were *κ* = 5.0, *ε*_*w*_ = 78.5, electrostatic cutoff 12 Å, Lennard–Jones cutoff 9.5 Å (switching 8.0–9.5 Å), hydrogen-bond heavy-atom prefilter 3.5 Å, and Amber scaling factors SCEE = 1.2 and SCNB = 2.0. The final 1 *µ*s of each trajectory (10 000 frames saved every 0.1 ns) was analyzed after discarding the first and last 100 frames. The remaining frames (100–9900) were subsampled every 20 frames, corresponding to a 2 ns sampling interval (491 frames per trajectory).

