## Supplementary Information for "Rewiring V-type and K-type enzyme allostery through subunit interface mutations"

#### This PDF file includes:

Supporting text
Figs. S1 to S4
Table S1
SI References

### Supporting Information Text

### Materials and Methods

**Additional details on experimental setup.** . Both in the NMR and in the kinetic experiments, IGP was used in place of PRFAR because PRFAR binding causes pronounced line broadening and disappearance of HSQC resonances as a result of enhanced conformational exchange (1)

**Molecular dynamics simulations.** Structural models for the apo and holo forms of IGPS were constructed from the crystal structure of *Thermotoga maritima* IGPS (PDB ID: 1GPW, 2.4 Å resolution). In this structure, chain C displays loop 1 in a conformation compatible with effector binding; therefore, the apo model was generated by extracting chains C and D of the HisF–HisH complex and reverting the engineered mutation D11N to the wild-type residue. The PRFAR-bound structure was constructed following the protocol described previously. For the substrate-bound state, the glutamine position was determined by aligning HisH domains of the 7AC8 structure to 1GPW and placing glutamine in the pocket, followed by manual refinement to remove clashes.

Systems were parameterized using the Amber ff19sb force field together with the GAFF for the ligand. All crystallographic waters associated with the complex were retained. The system was solvated in an explicit water box of approximately $110 \times 110 \times 110$ Å, ensuring a minimum distance of 10 Å between the protein and the box boundaries, corresponding to roughly 20,000 water molecules. Counterions ($Na^+$ and $Cl^-$) 0.1 M was added until the system was neutralized.

Molecular dynamics simulations were carried out using the AMBER GPU implementation. Following energy minimization and equilibration (see Supplementary Information for details), production trajectories were generated for both apo and PRFAR-bound systems.

To compare the effects of effector binding and mutants, simulations were performed for apo enzyme, substrate- and PRFAR-bound states and ternary state with both gln and PRFAR bound in the respective pockets. These systems are referred to as E • apo, E • gln, E • pr, E • ter. Each of these was prepared both in the WT state and introducing the hR18A or hK181A (HisH) mutations. Each system was simulated for approximately 1.5 $\mu$s. The final 1 $\mu$s of each trajectory was used for analysis, from which ∼10,000 frames were extracted at regular intervals. To ensure statistical robustness, three independent replica simulations were performed for each condition.

We employed the following pre-equilibration procedure: minimization of the solvent, constraining the rest of the atoms at the crystal structure positions (100 ps). The optimized solvated complex was then slowly heated to 303 K (for 30°C simulations) for a minimum of 100 ps. Subsequent MD simulations (200 ps) in the canonical NVT ensemble were performed using Langevin dynamics, with applied harmonic constraints to the protein and PRFAR heavy atoms, with force constants set to 1 kcal/mol·Å$^2$. During this heating procedure, different parts of the system were gradually unconstrained until all atoms were set freed. Unconstrained MD simulations were run for more than 4 ns, for a total pre-equilibration simulation time of at least 5 ns. Finally, MD simulations were performed in the NPT ensemble at 303 K and 1 atm using the Langevin piston. The MD simulations were carried out for at least 1200 ns in three replicas for each state. All simulations were performed using periodic boundary conditions. Electrostatic interactions were treated with the Particle Mesh Ewald method, and van der Waals interactions were calculated using a switching distance of 10 Åand a cutoff of 12 Å. We used a multiple time-stepping algorithm to evaluate bonded, short-range non-bonded, and long-range electrostatic interactions at every one, two, and four timesteps, respectively, using a timestep of integration set to 2 fs. All subsequent trajectory analysis was carried on the last $1\mu$ of production (for each replica), sampled every 0.1 ns, between frame 100 and 9,900, using MDAnalysis and MDigest suite, where not specified otherwise.

**Traffic Analysis.** To characterize dynamical communication pathways within IGPS, we employed correlation networks derived from MD trajectories. In this framework, the protein is represented as a graph in which each node corresponds to the C$\alpha$ atom of a residue, and edges represent dynamical correlations between residue motions.

Edges are defined through an adjacency matrix $A$, whose elements are the generalized correlation coefficients $r_{MI}(x_i, x_j)$ between residues $i$ and $j$:

$$r_{MI}(x_i, x_j) = \left[1 - \exp\left(-\frac{2}{3} I(x_i, x_j)\right)\right]^{1/2},$$

where $I(x_i, x_j)$ is the mutual information between the positional fluctuations of the two residues:

$$I(x_i, x_j) = S(x_i) + S(x_j) - S(x_i, x_j).$$

Here $S(x_i)$ and $S(x_i, x_j)$ denote the marginal and joint Shannon entropies:

$$S(x_i) = -\int dx_i\, p(x_i) \ln p(x_i),$$

$$S(x_i, x_j) = -\iint dx_i\, dx_j\, p(x_i, x_j) \ln p(x_i, x_j),$$

where $p(x_i)$ and $p(x_i, x_j)$ are the probability distributions of atomic displacements sampled from the MD trajectories. The generalized correlation coefficient ranges from 0 (uncorrelated motions) to 1 (fully correlated motions).

 Apala Chaudhuri,[a,b,1] Federica Maschietto,[c,1] Olivia Enny,[a] Victor S. Batista,[a] J. Patrick Loria[a,d]

To identify the dominant traffic routes between the effector-binding pocket and the catalytic site, the network of dynamical correlations was treated as follows. The effective distance between residues $i$ and $j$ is defined as

$$w_{ij} = -\log\left(r_{MI}(x_i, x_j)\right),$$

so that strongly correlated residue pairs correspond to shorter distances. Minimizing the total path length,therefore, identifies routes that maximize the dynamical correlation between the starting and ending nodes.

In the present analysis, traffic maps were constructed from precomputed residue-residue generalized linear mutual information matrices obtained from concatenated MD trajectories for each system. For each variant, the mutual-information matrix $w$ was used as the weighted adjacency matrix of an undirected graph. Self-connections were removed by setting the diagonal elements to zero, and any non-finite entries were replaced by zero prior to graph construction. Edges corresponding to residues found at a distance $< 6$ Åin more than 75% of the sampled configurations are then used to build the corresponding protein network.

To quantify communication traffic through the network, we computed both edge betweenness centrality and node betweenness centrality on the weighted graph using the shortest path formalism implemented in NetworkX. Edge betweenness centrality measures the fraction of weighted shortest paths passing through a given edge, whereas node betweenness centrality measures the corresponding fraction passing through a given node. Both quantities were computed in normalized form. In practice, these metrics identify edges and residues that are most frequently traversed by the shortest communication routes and therefore act as dominant conduits of allosteric signal propagation. Because absolute centrality values differ across systems, edge and node betweenness values were rescaled independently within each network by min–max normalization to the interval $[0, 1]$: $\tilde{b}_k = \frac{b_k - b_{\min}}{b_{\max} - b_{\min}}$, with $\tilde{b}_k = 0$ if $b_{\max} = b_{\min}$. The resulting normalized edge and node centralities were used for downstream visualization.

For structural rendering, normalized edge betweenness values were mapped onto residue pairs and visualized in PyMOL as cylinders connecting C$\alpha$ atoms. Edge widths were scaled proportionally to the corresponding normalized traffic value. Node betweenness values were supplied independently and used to modulate the size of spheres shown at the corresponding C$\alpha$ positions. Residue coordinates were taken from a reference PDB structure, and only C$\alpha$ atoms were used to define node positions. For clarity, only edges with normalized edge betweenness above a threshold of 0.2 were displayed in the final traffic maps. Edge colors were assigned from a continuous colormap after independent normalization of the displayed edge values. The final traffic representations therefore encode two complementary observables: edge thickness reports the relative importance of residue-residue communication links, while sphere size reports the relative traffic concentrated at individual residues. These maps were used to compare how WT, hK181A, and hR18A redistribute dynamical communication across the HisF–HisH interface and through the allosteric network.

**Non-covalent interaction analysis and delta effective scores.** For each trajectory, we computed atom-level non-covalent interactions for every analyzed frame and subsequently aggregated them into residue pairs, independently of donor-acceptor orientation. For each residue pair, we first determine the percentage occupancy $p_{ij}$ as the fraction of frames in which a given atom-level interaction is present. All atom-level interactions connecting the same residue pair are then combined, and the number of distinct contributing atomic contacts $n_{ij}$ is recorded.

To account simultaneously for interaction persistence and multiplicity, we define an effective interaction score that weights the mean occupancy of the contributing contacts by their number (additional details in SI). Scores are computed independently for each mutant, conformational state, and trajectory replicate, and are subsequently averaged across the three 1 $\mu$simulation replicates.

Mutant contrasts ($\Delta$) are defined as the difference in effective scores between two mutants for each conformational state. For every residue pair, we report the mean $\Delta$ across the four states, while the whiskers correspond to the standard error computed from the distribution of these state-specific differences.

***Non-covalent interactions.*** We compute hydrogen bonds, hydrophobic contacts, and $\pi$–$\pi$ stacking interactions at the atom level for every analyzed frame of each trajectory. For hydrogen bonds, we evaluate all donor-hydrogen-acceptor triplets and assign a contact when the angle lies between 120° and 180° and the H$\cdots$A distance is $\leq 4.0$ Å.

For hydrophobic contacts, we identify hydrophobic heavy atoms (C, S, halogens), compute residue-level centers of mass for the interacting atoms, and assign a contact when the inter-residue distance is $\leq 3.6$ Å.

For $\pi$–$\pi$ stacking interactions, we compute ring centroids and normals for aromatic residues (Phe, Tyr, His, Trp), and assign a contact when the centroid distance is $\leq 4.2$ Å and the inter-ring angle falls within face-to-face (0°–45°) or edge-to-face (45°–135°) regimes.

***Residue-pair aggregation and effective scores.*** All atom-level interactions connecting residues $i$ and $j$ are grouped into a single unordered residue pair $(i, j)$, treating donor/acceptor orientation as equivalent, and exact self-contacts are discarded. For each pair, we compute the percentage occupancy $p_{ij}$ as the mean occupancy of the contributing atom-level contacts and record the number of distinct contacts $n_{ij}$. These quantities are combined into an effective interaction score,

$$S_{ij} = p_{ij}\left[1 + \Big(\ln(1 + n_{ij}) - \ln 2\Big)\right],$$

which equals $p_{ij}$ when a single atomic contact contributes ($n_{ij} = 1$) and increases when multiple contacts jointly support the interaction.

Effective scores are computed independently for each mutant, conformational state, and replicate, and then averaged across replicates. Mutant contrasts are defined as $\Delta = S_{ij}^{(A)} - S_{ij}^{(B)}$ for each state. For every residue pair, we report the mean $\Delta$ across the four states, while the whisker represents the standard error of these state-specific differences, capturing how consistently each mutation perturbs the corresponding interaction.

**Energy Decomposition Analysis.** We performed per-residue interaction energy decomposition on "dried" MD trajectories to localize energetic changes associated with mutation and effector binding. Pairwise atom–atom interaction energies were computed for each selected frame and subsequently summed to obtain per-residue contributions. The final 1 $\mu$s of each trajectory (10 000 frames saved every 0.1 ns) was analyzed after discarding the first and last 100 frames. The remaining frames (100–9900) were subsampled every 20 frames, corresponding to a 2 ns sampling interval (491 frames per trajectory). Each saved frame was analyzed using MDAnalysis for selections and ParmEd for topology and parameters. Energies were computed as follows.

***Electrostatics.*** Electrostatic interactions were computed on the dried trajectories using a distance-dependent dielectric model,(2) $\epsilon(r) = \kappa r$, which provides an approximate treatment of solvent screening in the absence of explicit solvent. The pairwise electrostatic interaction between atoms $i$ and $j$ is therefore

$$E_{\mathrm{ele}}(i,j) = \frac{K_e\, q_i q_j}{\kappa\, r_{ij}}, \qquad [1]$$

where $K_e = 332.0636$ kcal Å mol$^{-1}$ e$^{-2}$, $q_i$ and $q_j$ are the atomic charges, and $r_{ij}$ is the interatomic distance. The parameter $\kappa = 5.0$ controls the strength of dielectric screening and was chosen to represent moderate solvent attenuation while preserving physically meaningful electrostatic interactions within the protein interior.

***Van der Waals.*** Van der Waals interactions were computed using a Lennard–Jones potential,

$$E_{\mathrm{vdw}}(i,j) = S(r_{ij})\, 4\epsilon_{ij} \left[ \left( \frac{R_{\mathrm{min},ij}}{r_{ij}} \right)^{12} - \left( \frac{R_{\mathrm{min},ij}}{r_{ij}} \right)^{6} \right], \qquad [2]$$

where $\epsilon_{ij}$ and $R_{\mathrm{min},ij}$ are the standard Lennard–Jones parameters.

To ensure a smooth truncation of the potential near the cutoff, a quintic switching function was applied,

$$S(r) = 1 - 10t^3 + 15t^4 - 6t^5, \qquad t = \frac{r - r_s}{r_c - r_s}.$$

The switching function is applied for $r_s < r < r_c$, with $S(r) = 1$ for $r \leq r_s$ and $S(r) = 0$ for $r \geq r_c$. In this work we used $r_s = 8.0$ Å and $r_c = 9.5$ Å, values consistent with the non-bonded interaction settings used during the MD simulations.

***Hydrogen-bonds.*** Hydrogen-bond interactions were modeled using an orientation-dependent term,

$$E_{\mathrm{hb}} = -E_0\, s_r\, s_\theta, \qquad [3]$$

where $E_0 = 0.8$ kcal/mol sets the maximum strength of the hydrogen-bond interaction and $s_r$ and $s_\theta$ are distance- and angle-dependent switching functions. Hydrogen bonds were identified using geometric criteria ($\mathrm{H} \cdots \mathrm{A} \leq 2.5$ Å and donor–hydrogen–acceptor angle $\geq 140°$).The radial switching function $s_r$ is controlled by parameters $r_0 = 3.0$ Å and $k = 0.20$, where $r_0$ defines the characteristic donor–acceptor distance of the interaction and $k$ controls the steepness of the distance-dependent decay of the hydrogen-bond weight.

This term is intended to provide a geometry-weighted correction that emphasizes well-formed hydrogen bonds rather than a full physical hydrogen-bond potential, since electrostatic and van der Waals interactions are already accounted for in the base energy terms.

***Solvent corrections.*** To approximate solvent screening effects, additional polar and nonpolar solvation corrections were included. Polar corrections were estimated using a screened Coulomb interaction,

$$E_{\mathrm{polar,corr}}(i,j) = w_{ij}\, K_e\, q_i q_j \left( \frac{1}{\varepsilon_w} - \frac{1}{\kappa r_{ij}} \right) \frac{1}{r_{ij}}, \qquad [4]$$

where $\varepsilon_w = 78.5$ is the dielectric constant of water and $w_{ij}$ is a distance-dependent attenuation factor.

Nonpolar solvation contributions were approximated from the solvent-accessible surface area (SASA),(3)

$$E_{\mathrm{np}}(\mathrm{res}) = \gamma\, \mathrm{SASA}_{\mathrm{res}}, \qquad [5]$$

where $\mathrm{SASA}_{\mathrm{res}}$ is the solvent-accessible surface area of the residue and $\gamma = 0.006$ kcal mol$^{-1}$ Å$^{-2}$ is an empirical surface tension coefficient commonly used to approximate hydrophobic solvation effects.

 

***Composite terms.*** For each residue, total interaction energies were obtained by summing all pairwise atom–atom contributions involving atoms belonging to that residue. The resulting base interaction energy was defined as

$$E_{\mathrm{base}} = E_{\mathrm{ele}} + E_{\mathrm{vdw}} + E_{\mathrm{polar,corr}} + E_{\mathrm{np}}. \tag{6}$$

Energetic effects of mutation were quantified as

$$\Delta E(\mathrm{res}) = E^{\mathrm{mut}}_{\mathrm{base}}(\mathrm{res}) - E^{\mathrm{WT}}_{\mathrm{base}}(\mathrm{res}), \tag{7}$$

while mutation-induced changes in binding energetics were computed as

$$\Delta\Delta E(\mathrm{res}) = \left[E^{\mathrm{bound}} - E^{\mathrm{apo}}\right]_{\mathrm{mut}} - \left[E^{\mathrm{bound}} - E^{\mathrm{apo}}\right]_{\mathrm{WT}}. \tag{8}$$

Unless otherwise noted, the following parameters were used throughout the analysis: distance-dependent dielectric parameter $\kappa = 5.0$, water dielectric constant $\varepsilon_w = 78.5$, electrostatic cutoff 12 Å, Lennard–Jones cutoff 9.5 Å with switching between 8.0–9.5 Å, hydrogen-bond heavy-atom prefilter 3.5 Å, explicit $\mathrm{H}\cdots\mathrm{A} \leq 2.5$ Å and donor–hydrogen–acceptor angle $\geq 140^\circ$, Amber scaling factors SCEE = 1.2 and SCNB = 2.0, and nonpolar surface tension coefficient $\gamma = 0.006\ \mathrm{kcal\,mol^{-1}\,Å^{-2}}$.

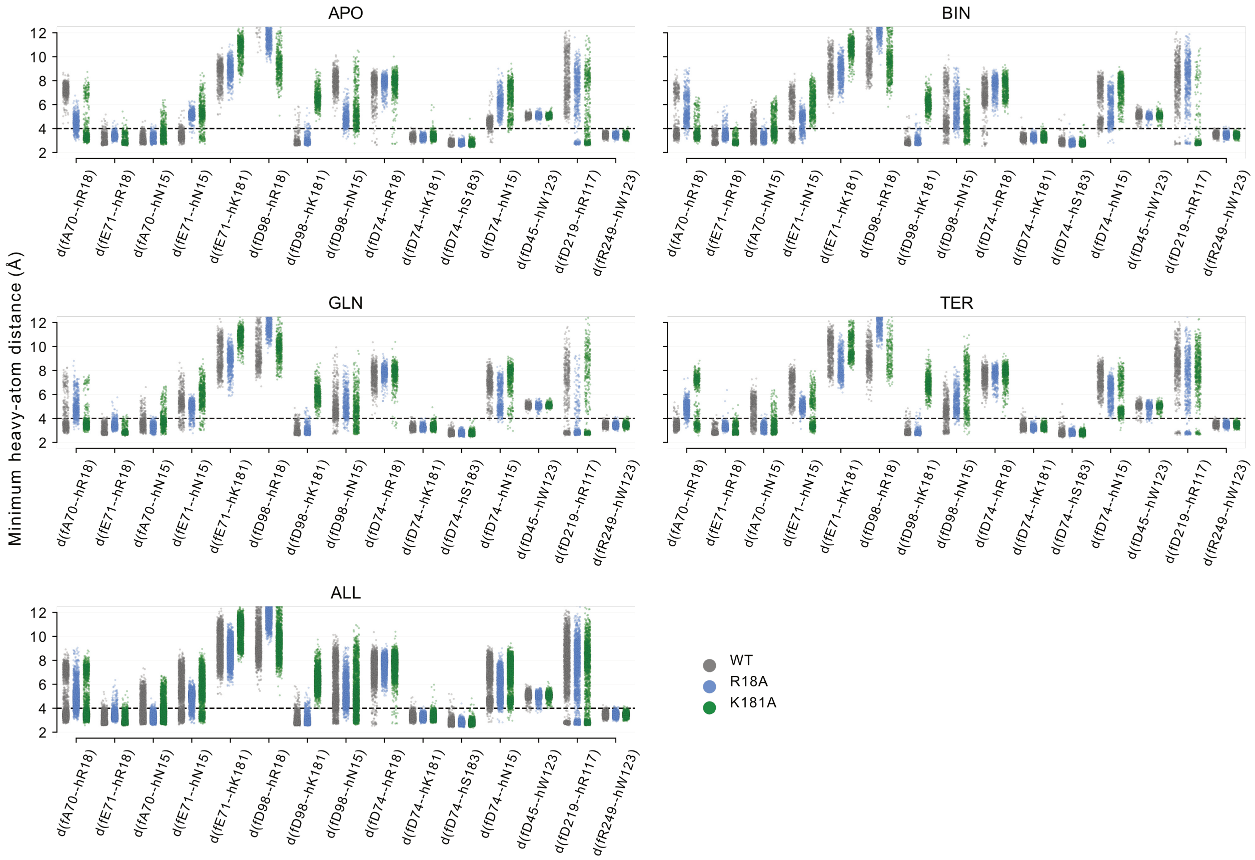


**Fig. S1.** Minimum heavy-atom distance distributions for selected HisF–HisH interfacial residue pairs across apo, binary PRFAR-bound, glutamine-bound, ternary PRFAR-bound, and pooled states. Each point reports the minimum heavy-atom distance between the two residues in one sampled MD frame. The ALL panel concatenates trajectories from all four ligand states for each system. Contacts are labeled as $d(\mathrm{residue\ pair})$ on the x-axis. The dashed horizontal line indicates the distance threshold used to define contact persistence. WT, R18A, and K181A are shown in gray, blue, and green, respectively.

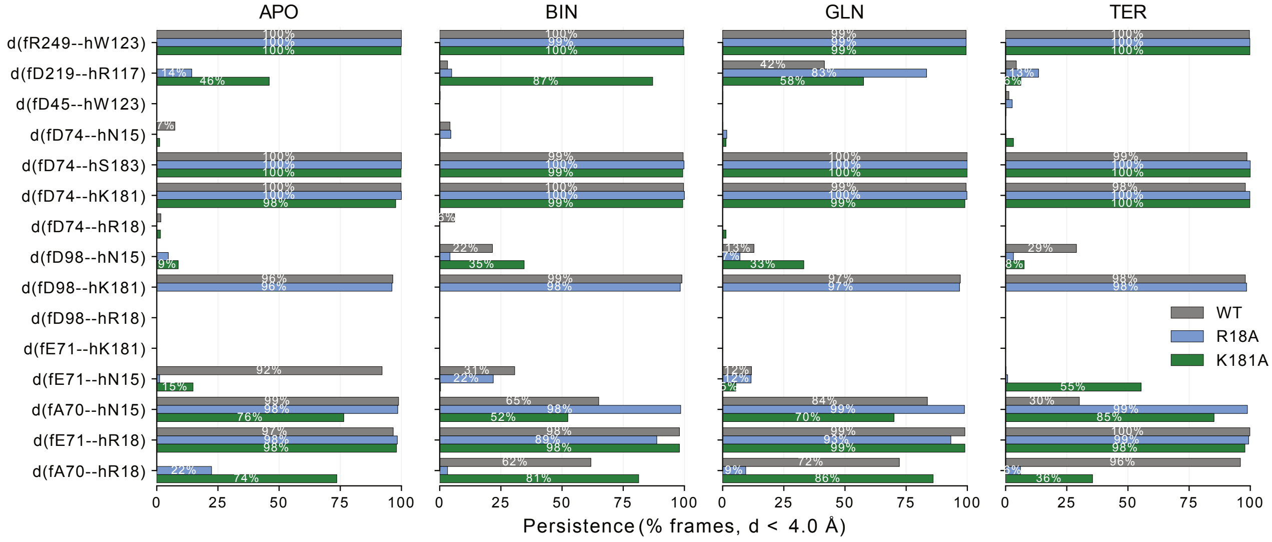


**Fig. S2.** State-resolved persistence of selected HisF–HisH interfacial contacts. Persistence is defined as the percentage of sampled MD frames in which the minimum heavy-atom distance between the two residues is below the contact threshold. Bars report the mean persistence across three independent replicas. Contact pairs are shown for apo, binary PRFAR-bound, glutamine-bound, and ternary PRFAR-bound states. WT, R18A, and K181A are shown in gray, blue, and green, respectively.

 Apala Chaudhuri,[a,b,1] Federica Maschietto,[c,1] Olivia Enny,[a] Victor S. Batista,[a] J. Patrick Loria[a,d]

**Table S1. Glutamine-dependent activity measurements for WT, K181A, and R18A IGPS. Changes in absorbance/second (Abs/s). Values are reported as mean ± SD from triplicate measurements at 505 nm. Missing measurements are indicated by –.**

| Variant | [Gln] (mM) | Basal Δ Abs/s | Basal $Abs_{505}$ |
|---|---|---|---|
| WT | 0.1 | – | $2.10 \times 10^{-5} \pm 2.42 \times 10^{-5}$ |
| WT | 0.2 | $0.0013 \pm 5.12 \times 10^{-4}$ | $4.72 \times 10^{-5} \pm 2.05 \times 10^{-5}$ |
| WT | 0.5 | $0.00364 \pm 2.42 \times 10^{-4}$ | $1.21 \times 10^{-4} \pm 2.90 \times 10^{-5}$ |
| WT | 1 | $0.00499 \pm 2.58 \times 10^{-4}$ | $1.87 \times 10^{-4} \pm 1.38 \times 10^{-5}$ |
| WT | 2 | $0.00742 \pm 5.03 \times 10^{-4}$ | $2.09 \times 10^{-4} \pm 1.42 \times 10^{-5}$ |
| WT | 4 | $0.00811 \pm 4.05 \times 10^{-4}$ | $2.68 \times 10^{-4} \pm 1.31 \times 10^{-5}$ |
| WT | 8 | $0.012 \pm 3.37 \times 10^{-4}$ | $2.64 \times 10^{-4} \pm 2.16 \times 10^{-5}$ |
| WT | 10 | – | $2.57 \times 10^{-4} \pm 2.08 \times 10^{-5}$ |
| WT | 20 | $0.0116 \pm 1.33 \times 10^{-4}$ | $2.30 \times 10^{-4} \pm 2.46 \times 10^{-5}$ |
| WT | 30 | $0.0119 \pm 0.00198$ | $1.78 \times 10^{-4} \pm 3.32 \times 10^{-5}$ |
| WT | 40 | $0.0114 \pm 9.51 \times 10^{-4}$ | $2.66 \times 10^{-4} \pm 2.11 \times 10^{-5}$ |
| WT | 50 | $0.0109 \pm 6.52 \times 10^{-4}$ | – |
| WT | 60 | $0.0116 \pm 8.30 \times 10^{-4}$ | $3.03 \times 10^{-4} \pm 2.43 \times 10^{-5}$ |
| K181A | 0.1 | – | – |
| K181A | 0.2 | $0.0378 \pm 8.83 \times 10^{-4}$ | $7.75 \times 10^{-4} \pm 8.62 \times 10^{-6}$ |
| K181A | 0.5 | $0.0723 \pm 8.90 \times 10^{-4}$ | $0.00203 \pm 3.28 \times 10^{-5}$ |
| K181A | 1 | $0.122 \pm 0.00395$ | $0.00463 \pm 1.30 \times 10^{-4}$ |
| K181A | 2 | $0.15 \pm 0.00245$ | $0.00799 \pm 1.70 \times 10^{-4}$ |
| K181A | 4 | $0.152 \pm 0.00471$ | $0.0125 \pm 2.36 \times 10^{-4}$ |
| K181A | 8 | $0.17 \pm 0.00226$ | $0.016 \pm 5.02 \times 10^{-4}$ |
| K181A | 10 | $0.192 \pm 0.00325$ | $0.0188 \pm 3.26 \times 10^{-4}$ |
| K181A | 20 | $0.187 \pm 0.00191$ | $0.0193 \pm 5.71 \times 10^{-4}$ |
| K181A | 30 | $0.177 \pm 0.00385$ | $0.0197 \pm 4.87 \times 10^{-4}$ |
| K181A | 40 | $0.181 \pm 0.00455$ | $0.0189 \pm 7.38 \times 10^{-4}$ |
| K181A | 50 | $0.18 \pm 0.00306$ | $0.0191 \pm 4.12 \times 10^{-4}$ |
| K181A | 60 | $0.162 \pm 0.00325$ | $0.0196 \pm 5.33 \times 10^{-4}$ |
| R18A | 0.1 | – | – |
| R18A | 0.2 | $6.49 \times 10^{-5} \pm 3.50 \times 10^{-5}$ | $1.22 \times 10^{-4} \pm 2.12 \times 10^{-4}$ |
| R18A | 0.5 | $7.70 \times 10^{-5} \pm 5.22 \times 10^{-6}$ | $7.37 \times 10^{-5} \pm 6.61 \times 10^{-5}$ |
| R18A | 1 | $1.86 \times 10^{-4} \pm 2.07 \times 10^{-5}$ | $6.41 \times 10^{-5} \pm 6.68 \times 10^{-5}$ |
| R18A | 2 | $3.53 \times 10^{-4} \pm 8.26 \times 10^{-6}$ | $2.02 \times 10^{-4} \pm 2.21 \times 10^{-5}$ |
| R18A | 4 | $4.45 \times 10^{-4} \pm 2.44 \times 10^{-5}$ | $3.02 \times 10^{-4} \pm 1.14 \times 10^{-5}$ |
| R18A | 8 | $5.19 \times 10^{-4} \pm 3.89 \times 10^{-5}$ | $4.16 \times 10^{-4} \pm 8.08 \times 10^{-6}$ |
| R18A | 10 | $5.00 \times 10^{-4} \pm 4.52 \times 10^{-6}$ | $4.74 \times 10^{-4} \pm 3.66 \times 10^{-5}$ |
| R18A | 20 | $5.95 \times 10^{-4} \pm 3.43 \times 10^{-5}$ | $4.48 \times 10^{-4} \pm 2.53 \times 10^{-5}$ |
| R18A | 30 | $5.19 \times 10^{-4} \pm 3.58 \times 10^{-5}$ | $5.24 \times 10^{-4} \pm 2.51 \times 10^{-5}$ |
| R18A | 40 | $4.78 \times 10^{-4} \pm 3.11 \times 10^{-5}$ | $5.45 \times 10^{-4} \pm 1.67 \times 10^{-5}$ |
| R18A | 50 | $5.63 \times 10^{-4} \pm 1.92 \times 10^{-5}$ | $5.39 \times 10^{-4} \pm 4.98 \times 10^{-5}$ |
| R18A | 60 | $4.66 \times 10^{-4} \pm 1.21 \times 10^{-5}$ | $5.10 \times 10^{-4} \pm 4.54 \times 10^{-5}$ |

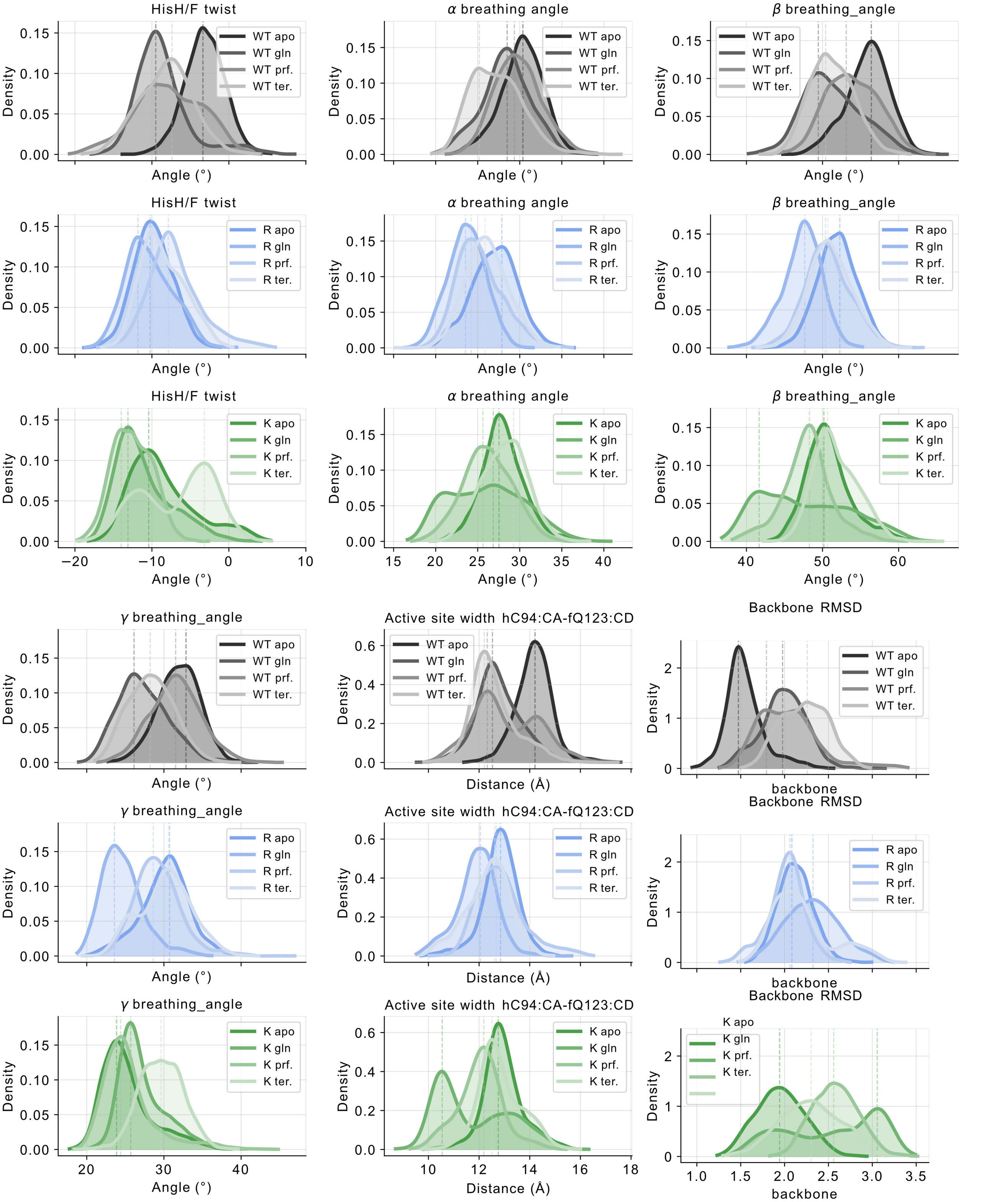


**Fig. S3.** Distributions of global and local dynamical metrics computed from MD trajectories of WT, R18A, and K181A IGPS across apo, glutamine-bound, PRFAR-bound, and ternary states. In addition to the HisF/H twist and the previously defined $\alpha$ breathing angle, the analysis includes related interdomain breathing coordinates: $\beta(fG166C_\alpha - hW/AzoF123C_\gamma - hE96C_\alpha)$, which monitors displacement of the HisH $\beta5$–$\beta6$ loop, and $\gamma(fL196C_\alpha - hAzoF123C_\gamma - hR117C_\alpha)$, which reports on the relative displacement of the opposing HisF $\alpha6$ helix and HisH $\beta6$–$\beta7$ loop.

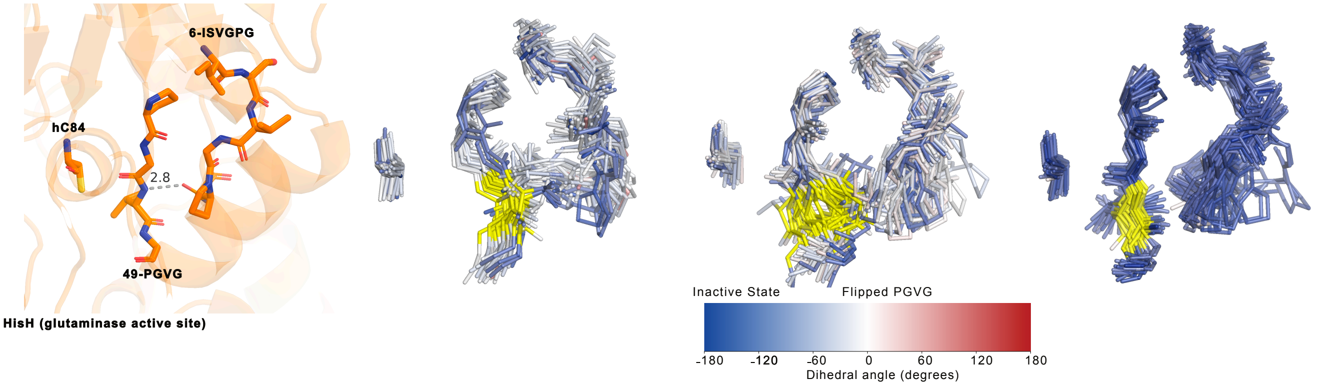


**Fig. S4.** Structural snapshots from selected binary and ternary MD trajectories illustrating the conformational sampling of the VAL304–GLY305 backbone dihedral in WT, K181A, and R18A IGPS. Only the local structural region surrounding residues 259–265, 302–305, and 337 is shown. Individual trajectory frames are split into separate conformers and colored according to the VAL304–GLY305 dihedral angle using a red–white–blue scale, where negative angles are red, values near 0° are white, and positive angles are blue. WT and K181A sample a broader range of dihedral states, spanning blue, white, and in some frames red conformations, whereas R18A remains predominantly confined to blue conformations. Because the transition toward white/red states corresponds to the local backbone flip associated with the active-like configuration, these snapshots indicate that WT and K181A are more prone than R18A to sample active-state conformations in the selected binary and ternary trajectories.
